# βTrCP is involved in the localization of the MRN complex on chromatin to enhance DNA damage repair

**DOI:** 10.1101/2025.03.06.640826

**Authors:** Alejandro Belmonte-Fernández, Joaquín Herrero-Ruíz, Carmen Sáez, Miguel Á. Japón, Mar Mora-Santos, Francisco Romero

## Abstract

Genomic instability underlies various diseases, including cancer. This instability arises from defects in critical cellular processes, particularly those involved in DNA damage repair. Therefore, a detailed understanding of these repair mechanisms is essential for developing strategies to prevent or diagnose such diseases. The MRN complex, composed of MRE11, NBS1, and RAD50, is among the earliest elements involved in detecting DNA damage. Upon detecting DNA breaks, this complex triggers a cascade of signaling events that regulate both cell cycle arrest and DNA repair. These signaling pathways are tightly controlled by various post-translational modifications, notably ubiquitination. Although several ubiquitin ligases have been implicated in different stages of the DNA damage response, our knowledge remains limited. In this study, we reveal that βTrCP, a substrate-recognizing subunit of the SCF (SKP1/CUL1/F-box protein) ubiquitin ligase, interacts *in vivo* with the proteins of the MRN complex. These interactions occur in normally proliferating cells and are dependent on the GSK3 kinase. Moreover, we show that βTrCP enhances the recruitment of the MRN complex to chromatin through MRE11, thereby promoting the efficient DNA damage repair. Hence, alterations in βTrCP function affecting MRN dynamics could have severe consequences for the cell homeostasis.

## INTRODUCTION

Double-strand breaks (DSBs) in DNA are the most deleterious DNA lesions for cells, and consequently, for the organism. In fact, if not properly repaired, they can lead to genomic instability, cellular senescence, chromosomal aberrations, immunodeficiency, and tumorigenesis ^1^. These breaks can be the consequence of cellular exposure to exogenous agents such as ionizing radiation or certain chemotherapeutic drugs, or they may arise from endogenous phenomena such as replicative stress or the generation of reactive oxygen species ^2,3^. Cells have two main pathways to repair DSBs: homologous recombination (HR) and non-homologous end joining (NHEJ), along with other secondary pathways ^4^. HR pathway repairs DNA without introducing errors, while NHEJ pathway is more prone to introduce mistakes. The MRN complex (MRE11, RAD50, NBS1) plays a crucial role in the DNA damage response. This heterohexameric complex is composed of two subunits of MRE11, two of RAD50, and two of NBS1. The catalytic core of the complex is formed by the two MRE11 subunits and the ATPase domains of RAD50, which together form the DNA-binding and processing area ^5^. NBS1, in contrast, acts as a scaffolding protein essential for interactions with other proteins ^6^. MRE11 interacts with both NBS1 and RAD50, while the latter two do not directly interact with each other to form the complex ^7^.

The MRN complex rapidly detects DNA damage and trigger the damage response cascade, including the arrest of the cell cycle ^8^. Furthermore, its nuclease activity, mediated by MRE11, contributes to both DNA end processing in NHEJ and end resection in HR ^9^. Additionally, the interaction between RAD50 subunits promotes the formation of bridges between the broken DNA molecule and its counterpart on the sister chromatid during HR. Similarly, it may also facilitate the alignment and approximation of broken DNA ends during NHEJ ^5^. In addition, the MRN complex is also involved in the repair of DNA-protein adducts, DSBs with forked ends, the processing of failed replication forks, telomere stability or detection, signaling, and defense against viral infections ^10-16^. DNA damage repair pathways are regulated by different types of post-translational modifications, such as phosphorylation, methylation, acetylation, and ubiquitination, among others ^2,17^. Specifically, ubiquitination plays several crucial roles in cells, ranging from protein degradation via the proteasome or lysosome to modifying the localization of ubiquitinated proteins, among other functions ^18^. The ubiquitination process involves several enzymes, including ubiquitin ligases, which catalyze the direct transfer of ubiquitin to the substrate protein. The SKP1/CUL1/F-box protein (SCF) ubiquitin ligase complex consists of several subunits, including F-box proteins, which are responsible for substrate recognition. The F-box βTrCP is involved in the ubiquitination of a wide variety of substrates associated with numerous cellular processes. βTrCP recognizes substrates through a specific binding motif known as degron, characterized by the canonical sequence D-S/T-G-X^2-3^-S/T (where X denotes any amino acid). Phosphorylation of this motif is typically required for recognition ^19,20^. Since βTrCP is involved in essential cellular processes, deregulation of its levels and/or activity has been linked to various diseases, including cancer ^21^. In this study, we identified and characterized the association between βTrCP and the proteins of the MRN complex. We found that βTrCP facilitates the localization of the MRN complex to chromatin, thereby promoting DNA damage repair.

## RESULTS

### βTrCP associates *in vivo* with the proteins of the MRN complex

In prior mass spectrometry analyses of HA βTrCP immunoprecipitates from transiently transfected Cos-7 cells ^22^, we identified some peptides corresponding to NBS1 and RAD50. To validate whether these proteins associate *in vivo* with HA βTrCP under physiological conditions, we performed individual immunoprecipitations of each protein followed by Western blot. As shown in Figures 1A and 1B, the immunocomplexes of NBS1, RAD50, and HA βTrCP each independently co-immunoprecipitated the other proteins. Given that MRE11 is a component of the MRN complex along with NBS1 and RAD50, we sought to examine its potential interaction with HA βTrCP. Our co-immunoprecipitation assays confirmed that MRE11 also interacts with the F-box protein, and that all these interactions take place specifically within the nuclear fraction of the cells (Figs. 1A, B). To rule out a potential effect of *HA βTrCP* overexpression due to transient transfection on the detected interactions, we aimed to verify that these interactions also occur at endogenous cellular levels. Figure 1C shows that endogenous βTrCP immunocomplexes also co-precipitated all the proteins of the MRN complex. In these experiments, we employed anti-β-catenin as a positive control for βTrCP immunoprecipitation ^23^, due to the lack of effective antibodies against βTrCP for Western blot assays. Additionally, using the SCF (βTrCP) complex produced and purified from insect cells, we demonstrated its ability to polyubiquitinate each of the MRN complex proteins separately, all of which were produced and labeled with [^35^S]-Met/Cys in reticulocyte extracts (Fig. 1D). These findings reveal that SCF (βTrCP) is a novel ubiquitin ligase for the MRN complex proteins.

**Figure 1.**
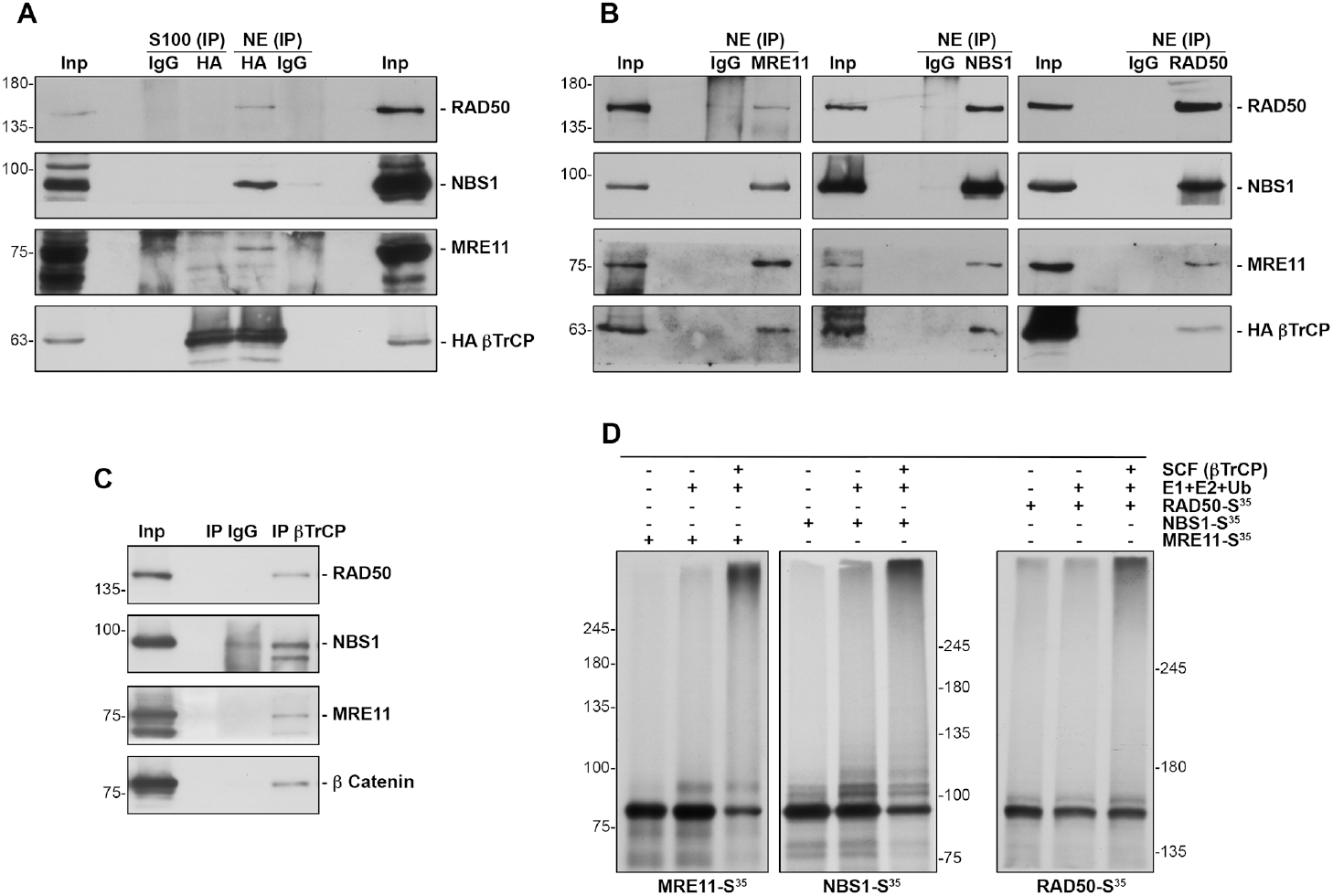
βTrCP interacts *in vivo* with the MRN complex proteins and induces their ubiquitination. **A**.Cytosolic (S100) and nuclear (NE) fractions from transiently transfected Cos-7 cells with pCS2HA βTrCP were used to immunoprecipitate HA βTrCP. Immunocomplexes were analyzed with the indicated antibodies. IgG: immunoprecipitation using normal mouse serum as a control. The input lane (Inp) was loaded with 1/20 of the extract used in each assay. **B**. Nuclear extracts (NE) from Cos-7 cells transfected with pCS2HA βTrCP were immunoprecipitated with anti-MRE11 (left panels), anti-NBS1 (central panels) or anti-RAD50 (right panels) and analyzed by Western blot. Immunoprecipitations using normal mouse serum (IgG) were used as controls. Inp: input of nuclear extracts. **C**. Similar to A and B, but using NP40 extracts from non-transfected Cos-7 cells. The immunoprecipitation efficiency of βTrCP was tested with anti-β catenin, a known partner of this F-box protein. **D**. Ubiquitination assay of *in vitro* transcribed and translated MRE11, NBS1 or RAD50 labeled with [^35^S] was carried out in the presence or absence of the following products: recombinant SCF (βTrCP) complex produced in Sf21 insect cells, E1 (His_6_-E1), E2 (His_6_-UbcH3 and UbcH5a), and ubiquitin (Ub). The samples were incubated at 30 °C for 1 hour and analyzed by SDS-PAGE and autoradiography. Polyubiquitylated proteins appear as a smear.

### MRE11 directly interacts with βTrCP

To identify the βTrCP binding site within each of the proteins of the MRN complex, we conducted an *in silico* study searching for potential consensus binding motifs for this F-box protein. Both MRE11 and NBS1 appear to harbor degrons that conform to the consensus motif D-S/T-G-X^2-3^-S/T, whereas RAD50 lacks such motifs. However, this does not exclude the possibility that RAD50 contains non-canonical motifs ^24^. Consequently, we introduced targeted mutations in the putative degrons of MRE11 and NBS1, as well as in a non-canonical site in RAD50 that most closely resembles the consensus motif ^25^. Using site-directed mutagenesis, we introduced point mutations by substituting serine or threonine, as well as acidic residues (which can mimic phosphorylated amino acids), with glycine or alanine (Fig. 2A). We then transfected cells with plasmids encoding either the wild-type or mutant versions of each protein and assessed their interaction with βTrCP through co-immunoprecipitation assays. We found that only the MRE11 mutant (Flag MRE11Δβ) abrogated the association (Fig. 2B). In this case, we used anti-PLK1 as a positive control for the βTrCP immunoprecipitation ^26^. This result implies that MRE11 is a direct target of βTrCP within the protein complex. Furthermore, this has also allowed us to localize the MRE11 degron between amino acids 596 and 603 of the protein. Nevertheless, it cannot be ruled out that NBS1 and/or RAD50 might also directly interact with βTrCP through other unidentified motifs or through the formation of MRN complexes with endogenous MRE11, as we will discuss later.

**Figure 2.**
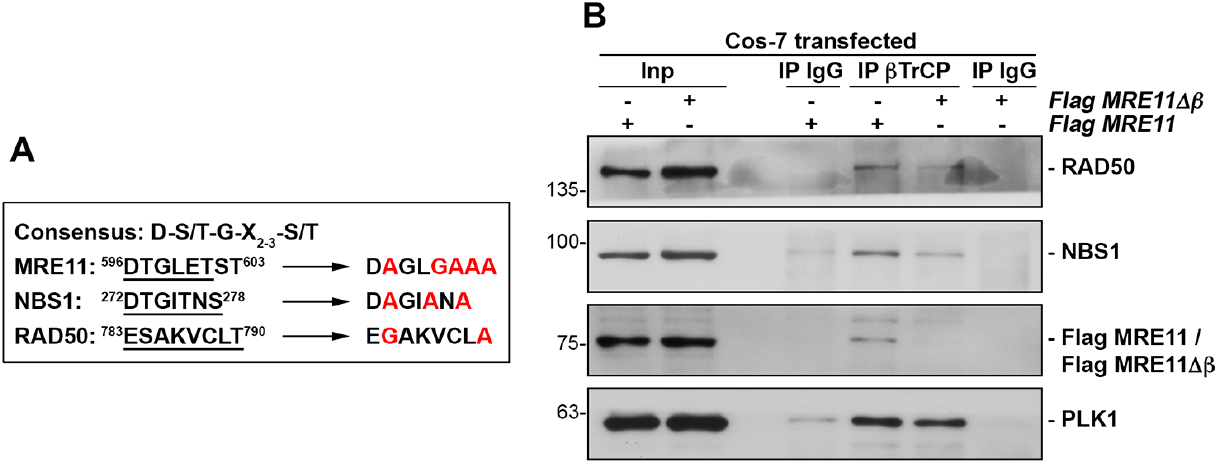
Identification of the βTrCP recognition motif in MRE11, whose mutation disrupts βTrCP/MRE11 interaction. **A**. *In silico* identification of potential consensus binding motifs for βTrCP in human MRE11 (residues 596 to 603), NBS1 (residues 272 to 278) and RAD50 (residues 783 to 790). Arrows indicate the amino acid changes (in red) introduced to generate mutants in these motifs. **B**. βTrCP was immunoprecipitated from NP40 extracts of Cos-7 transfected cells with pCMV6 Flag MRE11 or pCMV6 Flag MRE11Δβ, and coimmunoprecipitation of Flag MRE11 or Flag MRE11Δβ was assessed using anti-Flag. The presence of RAD50 and NBS1 in the immunocomplexes was also analyzed. Anti-PLK1 was used as a control of βTrCP immunoprecipitation. IP IgG: immunoprecipitation with normal rabbit serum as a control. Inp: NP40 extracts.

### GSK3 regulates the MRE11/βTrCP association

As mentioned in the *Introduction*, the association between F-box proteins and their substrates usually depends on prior phosphorylation of the degron motif in the substrate protein. The βTrCP motif in MRE11 includes threonine 597, whose phosphorylation by p70S6K has been previously established ^27^. Furthermore, using the Netphos 3.1 program, we found that serines or threonines 601, 602, and 603 could be potential targets of GSK3, a kinase traditionally involved in the association of substrates to βTrCP ^28^. To investigate these potential phosphorylation events, we treated cells with PF4708671, a p70S6K inhibitor ^29^, or lithium chloride, a GSK3 inhibitor ^30^. Interestingly, PF4708671 treatment did not substantially reduce the interaction between MRE11 and HA βTrCP, but primarily affected the RAD50/HA βTrCP association (Fig. 3A). As a control for the inhibitor, we examined the phosphorylation of threonine 389 in p70S6K, which occurs when its kinase activity is inhibited ^29^. However, lithium chloride treatment abrogated the interaction of all three MRN complex proteins with HA βTrCP (Fig. 3B). Similar results were obtained using CHIR99021, another GSK3 inhibitor ^31^, which was more effective at inhibiting the MRE11/βTrCP interaction than the NBS1/βTrCP or RAD50/βTrCP interactions (Fig. 3C). As a control, we analyzed the β catenin/βTrCP and WEE1/βTrCP associations, the former being GSK3β-dependent and the latter independent of GSK3β ^23,32^. As expected, the figures show that GSK3 inhibition reduced the β catenin/βTrCP interaction but did not affect the WEE1/βTrCP interaction. These results lead us to conclude that, in normally proliferating cells, p70S6K and, especially, GSK3 kinases are involved in the MRN/βTrCP association. Furthermore, since CHIR99021 treatment disrupted the MRE11/βTrCP interaction but partially preserved the associations with NBS1 and RAD50, likely due to its greater sensitivity to this inhibitor, it can be deduced that these latter associations are not mediated by MRE11.

**Figure 3.**
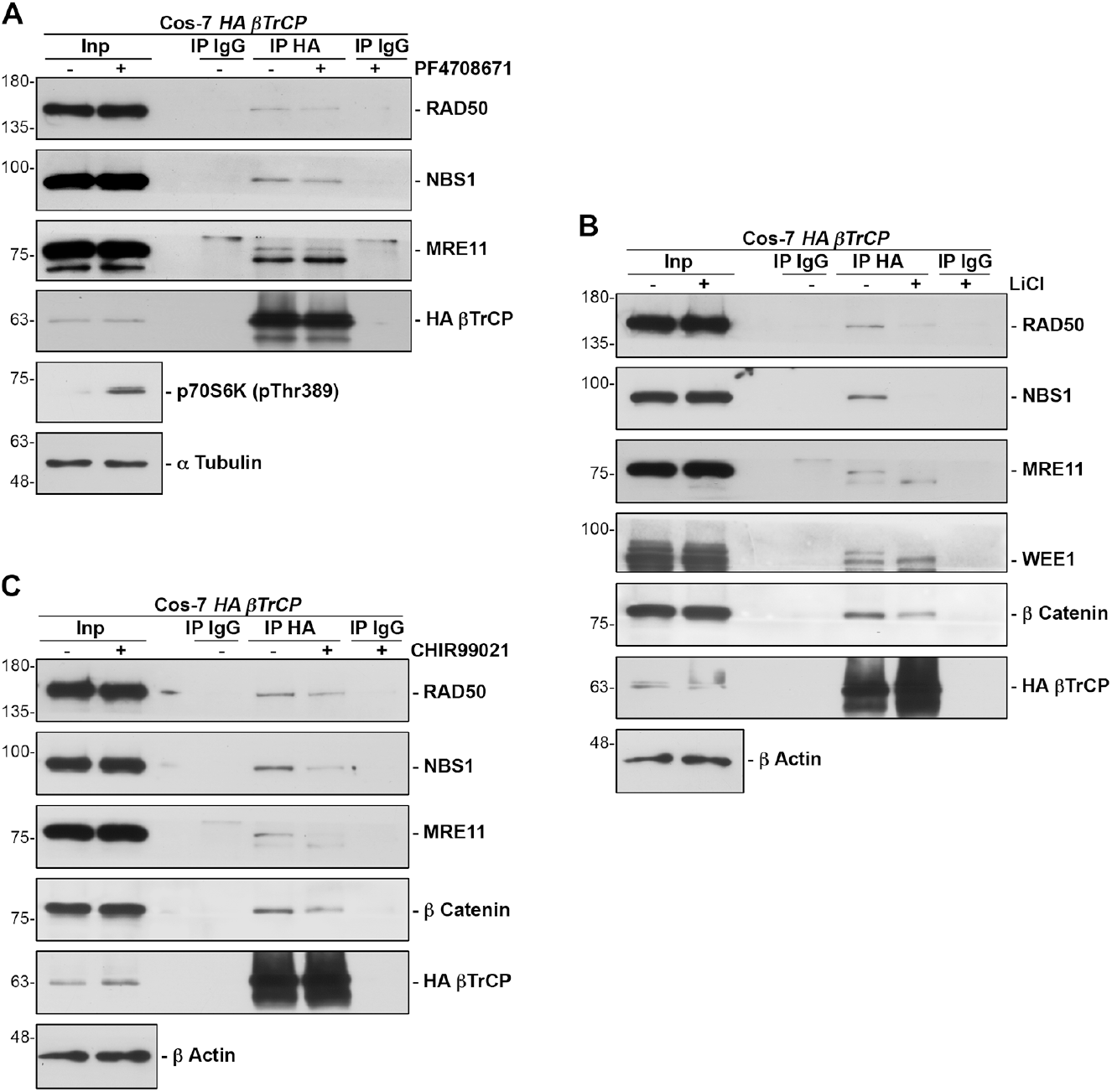
Involvement of p70S6K and GSK3 in the MRN/βTrCP interaction. **A**. Cos-7 cells were transfected with pCS2HA βTrCP and treated with PF4708671 (a p70S6K inhibitor) 18 hours prior to harvesting. NP40 extracts were prepared, and co-immunoprecipitation experiments were performed using an anti-HA antibody or normal mouse serum as control. Membranes were probed with the indicated antibodies. Anti-p70S6K (pThr389) was used to assess inhibitor efficiency. **B**. Similar to A, but cells were treated with lithium chloride, a GSK3 inhibitor. WEE1 and β catenin were used as controls, as described in the text. **C**. Similar to B, but cells were treated with CHIR99021, another GSK3 inhibitor. In this case, β catenin was analyzed as a control.

### βTrCP promotes the localization of MRN to chromatin

To understand the physiological role of the detected interaction, we first investigated whether the SCF (βTrCP) ubiquitin ligase is involved in the stability of the MRN complex proteins. To achieve this, we compared the levels of the three proteins in wild-type cells and in cells overexpressing *βTrCP*. However, no significant changes were observed in either the transiently or stably *βTrCP*-overexpressing cells (data not shown). We then investigated whether SCF (βTrCP) could affect MRN localization. For this, we used stable cell lines overexpressing *βTrCP, FBXW7* or their derivatives, from lentiviral constructs ^33^. As shown in the figure, *βTrCP* overexpression, but not *FBXW7*, which encodes another F-box protein that also associates with the MRN complex ^33^, enhances the localization of MRE11, RAD50, and NBS1 to chromatin (Fig. 4A). Moreover, this increase in MRN proteins on chromatin was prevented by overexpression of *βTrCPΔF*, which encodes a βTrCP mutant that binds to substrates but not to the rest of the ligase complex, thereby avoiding their ubiquitination (Fig. 4B). These results suggest that MRN polyubiquitination by SCF (βTrCP) facilitates its chromatin localization in unstimulated cells, likely promoting the repair of basal replication-associated DNA damage ^34^.

**Figure 4.**
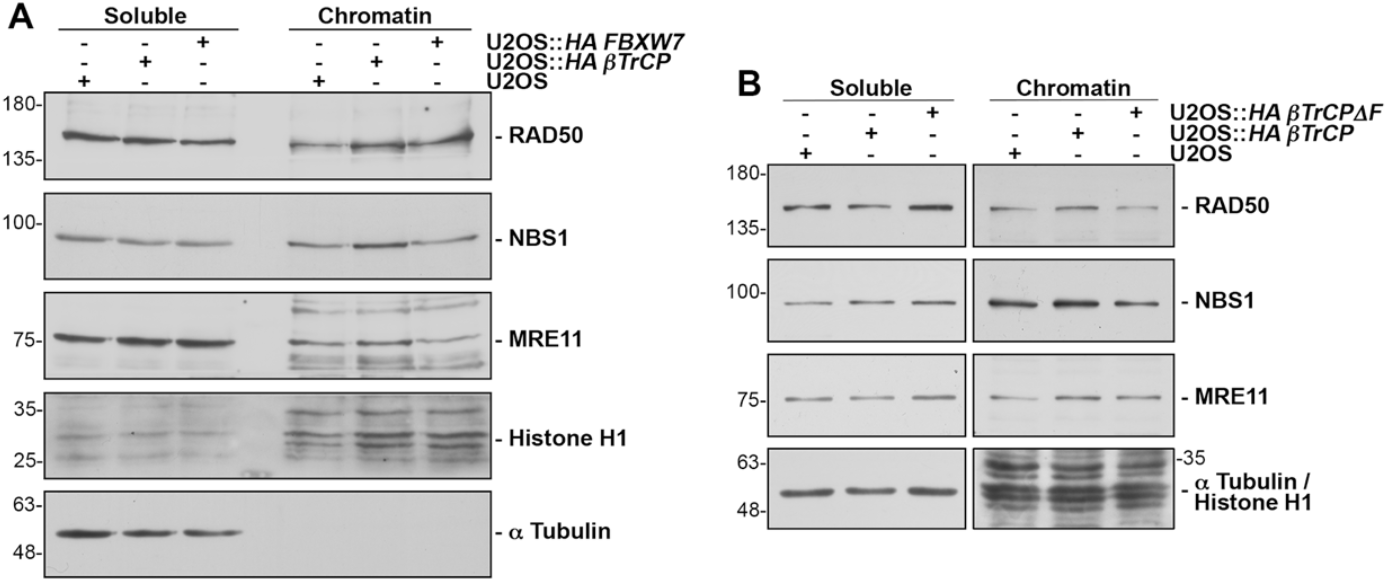
HA βTrCP enhances the accumulation of the MRN complex on chromatin. **A**. Soluble and chromatin fractions of U2OS cells and their lentiviral derivatives were analyzed by Western blotting. Fraction purity was verified by assessing the labeling of α tubulin and histone H1. **B**. Similar to A, but comparing the results obtained with the expression of the dominant-negative mutant HA βTrCPΔF.

### βTrCP stimulates the repair of ionizing radiation-induced DNA damage by recruiting the MRN complex to chromatin

To verify the hypothesis stated above, we expanded the study by inducing DNA damage using ionizing radiation. For this purpose, we constructed an inducible cell line expressing a dominant-negative mutant version of *βTrCP* (U2OS Flp-IN HA βTrCPΔF). Subsequently, we expressed or not the transgene for 24 hours, exposed the cells to X-ray radiation, and harvested at different time points. DNA damage was monitored by detecting phosphorylated H2AX histone (γH2AX) by Western blot. As shown in Figures 5A and 5B, in the absence of βTrCPΔF, H2AX phosphorylation decreases progressively over time, whereas its decrease is significantly delayed when *βTrCPΔF* is expressed. To investigate whether this repair delay caused by *βTrCPΔF* expression involves MRE11, we transfected U2OS cells with either pCMV6 Flag MRE11Δβ or the empty vector and examined the decrease in γH2AX over time following radiation exposure. Figure 5C shows that the MRE11 mutant, which cannot bind to βTrCP, similarly delays the reduction of H2AX phosphorylation. Additionally, Figure 5D shows, using fluorescence microscopy, that *βTrCPΔF* expression indeed reduces the localization of MRE11 to chromatin after irradiation. These findings indicate that MRE11 mediates the effect of βTrCP on DNA repair. Lastly, to quantify DNA damage at the single-cell level, we performed a comet assay, which efficiently measures DNA strand breaks. We compared comet formation in U2OS Flp-IN HA βTrCPΔF cells with and without transgene induction. As shown in Figure 5E, the overexpression of *βTrCPΔF* results in increased comet tail lengths, indicating an increase in DNA breaks. Collectively, these results allow us to conclude that the disruption of the βTrCP/MRE11 interaction delays DNA damage repair. Thus, βTrCP promotes repair by facilitating the recruitment of the MRN complex to chromatin.

**Figure 5.**
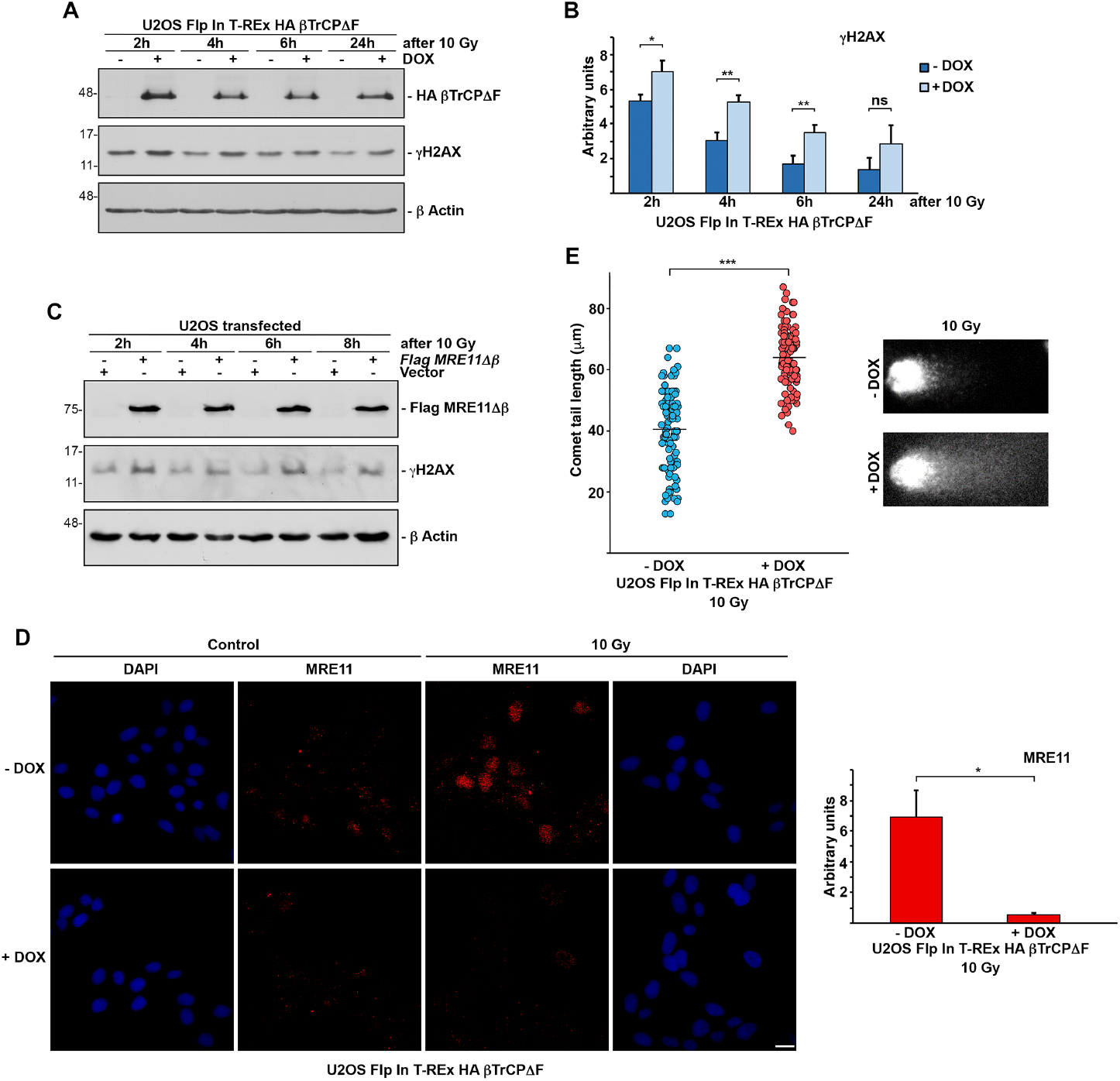
βTrCP promotes DNA damage repair through its association with MRE11. **A**. U2OS Flp In T-REx HA βTrCPΔF cells, induced or not with doxycycline (DOX), were irradiated (10 Gy) and harvested 2, 4, 6 or 24 hours later. Western blot analysis shows DNA damage repair, indicated by the decrease in γH2AX levels. **B**. The graph shows the quantification of γH2AX levels from Figure 5A, analyzed using ImageJ software. Error bars represent the standard deviation (SD) (n = 3). ns: non-significant, *p < 0.05, **p < 0.01 (Student’s *t* test). **C**. U2OS cells transfected with pCMV6 Flag MRE11Δβ or empty vector were irradiated (10 Gy) and harvested at the indicated times. As in A, γH2AX were used to assess DNA damage repair. **D**. U2OS Flp In T-REx HA βTrCPΔF cells were cultured directly onto pre-sterilized glass coverslips in the presence or absence of doxycycline (DOX), irradiated (10 Gy) or not, and treated 6 hours later with CSK buffer for 10 min to remove soluble fractions. The insoluble chromatin-associated fraction was analyzed by light microscopy. Scale bars represent 10 μm. The graph on the right shows the quantification of MRE11 levels in chromatin in cells expressing or not *HA βTrCPΔF*. *p < 0.05 (Student’s *t* test). **E**. U2OS Flp In T-REx HA βTrCPΔF cells, induced or not with doxycycline (DOX), were irradiated (10 Gy), and analyzed 1 hour later for DNA damage using the comet assay. Each dot represents the comet-tail length of an individual cell (at least 50 cells per condition). The line represents the average tail length for each condition. ***p < 0.001 (Student’s *t*-test). Representative comet images are shown on the right.

## DISCUSSION

Protein ubiquitination is a type of post-translational modification involved in many cellular processes ^35^. In particular, it plays a critical role in signaling cascades triggered by DNA damage. Upon DNA breaks, the MRN complex is recruited to damaged sites by γH2AX and RAD17 ^5^. γH2AX, facilitated by MDC1, interacts with NBS1, while RAD17, phosphorylated by ATM, directly associates with NBS1 ^36,37^. In this context, the ubiquitin ligases PELI1 and SKP2 generate Lys63-linked polyubiquitin chains on NBS1, promoting stable complex maintenance and facilitating further ATM activation ^38-40^. This results in a positive feedback loop that enhances ATM-related signaling. Subsequently, the ligases RNF8 and RNF168 are recruited to DSB sites and ubiquitinate histone H2A, which in turn promotes the recruitment of other key repair proteins, including 53BP1 and BRCA1 ^41,42^. These latter proteins are essential for choosing the repair pathway, either through HR or NHEJ. Consequently, they act antagonistically, initiating distinct and complex signaling cascades regulated by ubiquitination ^43^.

In this study, we identified SCF(βTrCP) as a novel ubiquitin ligase targeting the MRN complex proteins. We demonstrate both the *in vivo* association of βTrCP with MRE11, NBS1, and RAD50, as well as the direct ubiquitination of these three proteins by SCF(βTrCP). We further mapped the phosphodegron within MRE11, which directly binds to βTrCP following phosphorylation by GSK3. Additionally, we found that βTrCP facilitates the localization of MRE11 to chromatin following irradiation, thereby enhancing DNA damage repair. This result was corroborated by immunofluorescence and experiments using an MRE11 mutant unable to bind βTrCP. Therefore, βTrCP, along with the previously mentioned ligases PELI1 and SKP2, promotes the localization and/or stabilization of the MRN complex on chromatin. However, unlike SKP2 ^38^, βTrCP associates with MRN even under basal conditions. SKP2 is involved in the recruitment of ATM to chromatin upon genotoxic stress through the ubiquitination of NBS1, which would already be part of the MRN complex on chromatin ^38^. In contrast, our results show that βTrCP enhances the localization of MRN on chromatin, suggesting that it acts earlier than SKP2. PELI1, like SKP2, also contributes to the accumulation of DNA damage response proteins ^40^. PELI1 is primarily located in the cytoplasm of undamaged cells and translocates to DNA break sites upon ionizing radiation exposure. Nevertheless, it has been reported that PELI1 interacts with the MRN complex under basal conditions and that this interaction increases following ionizing radiation ^40^. However, the βTrCP/MRN interaction does not undergo substantial changes following DNA damage induction (data not shown), suggesting an earlier involvement in MRN complex function/localization. Additionally, PELI1 appears to participate in the proteasomal degradation of MRN complex proteins, whereas βTrCP does not.

As previously noted, our findings demonstrate that βTrCP has a positive effect on DNA damage repair, helping to prevent genetic information loss due to mutations or severe chromosomal rearrangements, which could ultimately lead to cancer development. Moreover, βTrCP also promotes cell cycle arrest, allowing for the effective processing and repair of DNA lesions. The activation of CHK1 and CHK2 triggers the phosphorylation of CDC25, facilitating its ubiquitination by the SCF(βTrCP) complex and subsequent degradation, leading to a reduction in active cyclin-CDK complexes ^44,45^. Additionally, SCF(βTrCP) is also involved in the degradation of MDM2, a negative regulator of p53, thereby supporting the accumulation of p53 and consequent cell cycle arrest through the induction of p21 CIP1 ^46^. Furthermore, SCF(βTrCP) also contributes to cell cycle arrest following UV-induced damage, facilitating repair by promoting the degradation of securin ^25^. Finally, once the damage has been repaired, SCF(βTrCP) induces the degradation of specific substrates such as WEE1 and claspin, which are involved in the ATR/CHK1 pathway, promoting cell cycle progression after damage repair ^47,48^. Consequently, βTrCP may act as a tumor suppressor in response to DNA damage. Accordingly, dysregulation of the levels and/or activity of this F-box protein has been linked to various types of cancer ^49-52^. Hypothetically, these cancers might be susceptible to specific DNA-damaging drugs. However, this might not always be the case, as cells can develop mechanisms allowing tumor cells to survive ^53,54^. Therefore, a deeper understanding of how the ubiquitination of specific proteins is implicated in damage repair may allow the development of more effective therapies for cancer treatment.

## MATERIALS AND METHODS

### Plasmids, cloning, point mutations, and sequencing

pCS2HA βTrCP, pCMVHA RAD50, pCMVHA NBS1, and empty vectors have been previously described ^33,55^. The pCMV6 Flag MRE11 was obtained from Origene (Rockville, MD, USA). To construct pLenti 6.3 HA βTrCPΔF and pCDNA5 FRT/TO HA βTrCPΔF, the corresponding PCR fragment was cloned into pLenti 6.3 or pCDNA5 FRT/TO, from Addgene (Watertown, MA, USA). The pCMV6 Flag MRE11Δβ mutant (MRE11 Thr597 to Ala, Glu600 to Gly, Thr601 to Ala, Ser602 to Ala, and Thr603 to Ala) was generated using the Q5 Site-Directed Mutagenesis Kit from New England Biolabs (Ipswich, MA, USA). Construct sequences and point mutations were verified by sequencing both strands with an automated sequencer.

### Cell line construction, cell culture, transfections, and drug treatments

A U2OS cell line with inducible expression of *HA βTrCPΔF* was constructed as previously described, using the pCDNA5 FRT/TO HA βTrCPΔF vector and the U2OS Flp In T-REx host cell line ^33^. The U2OS::*HA βTrCPΔF* cell line was obtained by retroviral infection using a retrovirus expressing *HA βTrCPΔF*, as described ^33^. U2OS and Cos-7 cells (from ATCC), as well as U2OS::*HA βTrCP*, U2OS::*HA FBXW7* ^33^, and the new cell lines U2OS::*HA βTrCPΔF* and U2OS Flp In T-REx HA βTrCPΔF were routinely cultured in Dulbecco’s Modified Eagle Medium (DMEM) from BioWest (Nuaille, France), as previously described ^56^. Plasmids were transiently transfected either by electroporation or using a lipid transfection reagent (Xfect; Takara Bio, Kusatsu, Japan) following the manufacturer’s standard instructions. Cells were harvested and lysed 18- or 48-hours post-transfection, respectively. In the indicated experiments, cells were treated with PF4708671 (10 μM, 18 hours, Tocris Bioscience, Bristol, UK), CHIR99021 (10 μM, 18 hours, Selleck Chemicals, Houston, TX, USA), LiCl (50 mM, 18 hours, Sigma-Aldrich, St. Louis, MO, USA), or doxycycline (DOX, 2 μg/mL, 24 hours, Sigma-Aldrich).

### Subcellular fractionation and cell lysis

Cytosolic (S100) and nuclear (NE) fractions were obtained as described ^57^. Soluble and chromatin fractions were separated using CSK buffer (10 mM HEPES (pH 7.4), 300 mM sucrose, 100 mM NaCl, 3 mM MgCl_2_, 0.5% Triton X-100, 1 μg/mL of each aprotinin, leupeptin and pepstatin, 1mM phenylmethylsulfonyl fluoride (PMSF)). After centrifugation at 5,000 *g* for 5 min at 4 °C, the soluble fractions were collected. The pellets were washed with the same lysis buffer and further solubilized in UTB buffer (8 M Urea, 50 mM Tris-HCl (pH 7.5), 150 mM β-Mercaptoethanol) and sonicated. Chromatin fractions were clarified by centrifugation at maximum speed for 30 min. Whole cell extracts were prepared at room temperature (RT) in UTB buffer (see above). The extracts were heated 5 minutes at 95 °C and centrifuged at 20,000 *g* for 20 min. NP40 extracts were prepared at 4 °C in 420 mM NaCl, 10 mM Tris-HCl (pH 7.5), 1% Nonidet P-40 (NP40), 10% glycerol, 1 mM PMSF, 1 μg/mL aprotinin, 1 μg/mL pepstatin, 1 μg/mL leupeptin and 10 μg/mL chymostatin for 20 min, centrifuged and then diluted to 150 mM NaCl. Extracts were frozen in liquid nitrogen and stored at -80 °C. Protein concentration was determined using the Bradford assay (Bio-Rad, Hercules, CA, USA).

### Electrophoresis, Western blot analysis, and antibodies

Proteins were separated by SDS-polyacrylamide gel electrophoresis (SDS-PAGE) and subsequently transferred onto nitrocellulose membranes (GE Healthcare, Little Chalfont, UK). The membranes were then incubated with the following antibodies: anti-HA-peroxidase monoclonal antibody (#12013819001, Roche, Basel, Switzerland); anti-β actin (sc-47778 HRP) and anti-MRE11 (sc-135992) monoclonal antibodies (Santa Cruz Biotechnology, Dallas, TX, USA); anti-α tubulin (T9026) and anti-Flag (F3165) monoclonal antibodies (Sigma-Aldrich); anti-PLK1 (#05-844) and anti-γH2AX (#05-636) monoclonal antibodies (EMD Millipore, Merck, Darmstadt, Germany); anti-β catenin (#610154) monoclonal antibody (BD Biosciences, Franklin Lakes, NJ, USA); anti-NBS1 (#14956), anti-WEE1 (#13084) and anti-phospho (Thr389) p70S6K (#9205) monoclonal antibodies (Cell Signaling Technology, Danvers, MA, USA); anti-histone H1 monoclonal antibody (ab11079, Abcam, Cambridge, UK); and anti-RAD50 monoclonal antibody (GTX70228, GeneTex, Irvine, CA, USA). Peroxidase-coupled donkey anti-rabbit IgG (NA934V) and sheep anti-mouse IgG (NA931) were acquired from GE Healthcare. Immunoreactive bands were visualized using the Enhanced Chemiluminescence (ECL) Western blotting system (GE Healthcare).

Protein level quantification was performed using ImageJ software (Image Processing and Analysis in Java; National Institutes of Health, Bethesda, MD, USA; http://imagej.nih.gov/).

### Co-immunoprecipitation experiments

1-2 mg of protein from S100 and NE fractions or NP40 extracts were incubated with normal mouse (sc-2025) or rabbit (sc-2027) sera (Santa Cruz Biotechnology) for 30 minutes, followed by incubation with protein G or A-Sepharose beads (GE Healthcare), respectively, for 1 hour at 4 °C. After centrifugation, the beads were discarded, and the supernatants were incubated for 2 hours with anti-HA (cat 901513, BioLegend, San Diego, CA, USA), anti-βTrCP (#4394, Cell Signaling Technology), anti-MRE11 (GTX70212, GeneTex), anti-NBS1 (#14956, Cell Signaling Technology), anti-RAD50 (NB110-60483, Novus Biologicals, Littleton, CO, USA), or normal mouse or rabbit sera, followed by incubation with protein G or A-Sepharose beads for 1 hour. The beads were washed, and the bound proteins were solubilized by adding SDS sample buffer and heated at 95 °C for 5 minutes.

### Irradiation procedure assays

Cells were exposed to different doses of ionizing radiation using a Philips MU 15F X-ray apparatus operated at 100 kV with a dose rate of 1 Gy/min. After specified time intervals, cells were harvested, and the extent of radiation-induced damage was assessed by the detection of γH2AX via Western blot analysis.

### Ubiquitination assays

Ubiquitinations of RAD50, NBS1 and MRE11 were performed in a volume of 10 μL containing 50 mM Tris-HCl (pH 7.6), 5 mM MgCl_2_, 0.6 mM DTT, 2 mM ATP, 2 μL of a recombinant SCF (βTrCP) complex produced in Sf21 insect cells, 1.5 ng/μL E1 (His_6_-E1; Boston Biochem, Cambridge, MA, USA), 10 ng/μL His_6_-UbcH3 (E2; Boston Biochem), 10 ng/μL UbcH5a (E2; Boston Biochem), 2.5 μg/μL ubiquitin (Millipore Sigma), 1 μM ubiquitin aldehyde (Boston Biochem), and 1 μL [^35^S]-Met/Cys-labeled *in vitro*-transcribed and -translated RAD50, NBS1 or MRE11 as substrates. The reactions were incubated at 30 °C for 1 hour and analyzed by SDS-PAGE and autoradiography.

### Comet assay

Comet assays were conducted following previously established protocols for alkaline unwinding and electrophoresis conditions ^58^. After 30 minutes in alkaline buffer, gel electrophoresis was carried out at 25 V for 30 minutes at 4 °C. Assays were performed on cells, with or without doxycycline treatment, 1 hour after exposure to 10 Gy or in the absence of radiation. Positive control cells were treated with 20 μM actinomycin D for 1 hour. Positive and negative controls were included with each experimental sample. A minimum of 50 individual cells per sample were analyzed in triplicate. Comet analysis was conducted using the ImageJ software.

### Immunofluorescence microscopy

Cells were cultured directly onto pre-sterilized glass coverslips in the presence or absence of doxycycline, irradiated (10 Gy) or not, and treated 6 hours later with CSK buffer for 10 min to remove soluble fractions. The insoluble cell fraction was fixed with 4% paraformaldehyde for 10 minutes on ice. Later, it was then washed twice with blocking buffer (1% BSA, 0.1% Tween 20) and incubated with the primary antibody for 1 h in the dark. After two PBS washes and one wash with the blocking buffer, it was incubated with the secondary antibody for 1 hours in the dark. Coverslips were subsequently washed three times with 1% PBS and mounted on microscopy slides using a mounting reagent with DAPI (Ibidi GmbH, Gräfelfing, Germany). Staining was analyzed using a Leica DMi8 inverted microscope with a 63x Plan Apo oil-immersion objective, under consistent laser parameters. All microscope images were analyzed with ImageJ software. The primary antibody used was anti-MRE11 mouse monoclonal antibody (GTX70212, GeneTex).

### Quantification and statistical analysis

Details regarding the statistical analyses, including the number of replicates and the use of standard deviation (SD), are provided in the figure legends. Statistical analyses were routinely performed with unpaired Student’s *t*-tests. Statistical significance is indicated in the figures as *p < 0.05, **p < 0.01 and ***p < 0.001.

## AUTHOR CONTRIBUTIONS

A.B.-F., J.H.-R. and F.R. performed study concept and design. A.B.-F., J.H.-R., and M.M.-S. provided acquisition, analysis and interpretation of data, and statistical analysis. C.S. and M.A.J. performed development of methodology. C.S., M.A.J. and F.R. wrote the manuscript with input from all other authors. A.B.-F. and J.H.-R. contributed equally to this work. All authors read and approved the final paper.

## ACKNOWLEDGEMENTS

This work was supported by grant PID2023-152257NB-C21 funded by MICIU/AEI/ 10.13039/501100011033/ ERDF, EU.

AB-F was the recipient of a Ph.D. fellowship from the Vicerrectorado de Investigaci6n Plan Propio de Investigaci6n (VI PPI) from Universidad de Sevilla. JH-R was a Talento Doctor fellow (Junta de Andalucia-FEDER). MM-S is a Marie Curie fellow (grant number 101024268).

## DATA AVAILABILITY

All data generated or analyzed during this study are included in this published article.

## REFERENCES

1. Scully, R., Panday, A., Elango, R. & Willis, N. A. DNA double-strand break repair-pathway choice in somatic mammalian cells. Nature Reviews Molecular Cell Biology vol. 20 698–714 Preprint at 10.1038/s41580-019-0152-0 (2019).

2. Cejka, P. & Symington, L. S. DNA End Resection: Mechanism and Control. Annual Review of Genetics vol. 55 285–307 Preprint at 10.1146/annurev-genet-071719-020312 (2021).

3. Aguilera, A. & Gómez-González, B. Genome instability: A mechanistic view of its causes and consequences. Nature Reviews Genetics vol. 9 204–217 Preprint at 10.1038/nrg2268 (2008).

4. Tan, J. et al. Double-strand DNA break repair: molecular mechanisms and therapeutic targets. MedComm vol. 4 e388 Preprint at 10.1002/mco2.388 (2023).

5. Bian, L., Meng, Y., Zhang, M. & Li, D. MRE11-RAD50-NBS1 complex alterations and DNA damage response: Implications for cancer treatment. Molecular Cancer vol. 18 1–14 Preprint at 10.1186/s12943-019-1100-5 (2019).

6. Rahman, S., Canny, M. D., Buschmann, T. A. & Latham, M. P. A Survey of Reported Disease-Related Mutations in the MRE11-RAD50-NBS1 Complex. Cells vol. 9 Preprint at 10.3390/cells9071678 (2020).

7. Lafrance-Vanasse, J., Williams, G. J. & Tainer, J. A. Envisioning the dynamics and flexibility of Mre11-Rad50-Nbs1 complex to decipher its roles in DNA replication and repair. Progress in Biophysics and Molecular Biology vol. 117 182–193 Preprint at 10.1016/j.pbiomolbio.2014.12.004 (2015).

8. Jin, M. H. & Oh, D. Y. ATM in DNA repair in cancer. Pharmacology and Therapeutics vol. 203 Preprint at 10.1016/j.pharmthera.2019.07.002 (2019).

9. Rupnik, A., Grenon, M. & Lowndes, N. The MRN complex. Current Biology vol. 18 Preprint at 10.1016/j.cub.2008.03.040 (2008).

10. Li, Y. et al. Nonhomologous end-joining with minimal sequence loss is promoted by the Mre11-Rad50-Nbs1-Ctp1 Complex in Schizosaccharomyces pombe. Genetics 206, 481–496 (2017).

11. Stracker, T. H., Carson, C. T. & Weilzman, M. D. Adenovirus oncoproteins inactivate the Mre11-Rad50-NBs1 DNA repair complex. Nature 418, 348–352 (2002).

12. Marcand, S. How do telomeres and NHEJ coexist? Mol Cell Oncol 1, (2014).

13. Marcomini, I. & Gasser, S. M. Nuclear organization in DNA end processing: Telomeres vs double-strand breaks. DNA Repair (Amst) 32, 134–140 (2015).

14. Aze, A., Zhou, J. C., Costa, A. & Costanzo, V. DNA replication and homologous recombination factors: Acting together to maintain genome stability. Chromosoma vol. 122 401–413 Preprint at 10.1007/s00412-013-0411-3 (2013).

15. Runge, K. W. & Li, Y. A curious new role for MRN in Schizosaccharomyces pombe non-homologous end-joining. Current Genetics vol. 64 359–364 Preprint at 10.1007/s00294-017-0760-1 (2018).

16. Deshpande, R. A., Lee, J. H., Arora, S. & Paull, T. T. Nbs1 Converts the Human Mre11/Rad50 Nuclease Complex into an Endo/Exonuclease Machine Specific for Protein-DNA Adducts. Mol Cell 64, 593–606 (2016).

17. Ray, U. & Raghavan, S. C. Understanding the DNA double-strand break repair and its therapeutic implications. DNA Repair (Amst) 106, (2021).

18. Damgaard, R. B. The ubiquitin system: from cell signalling to disease biology and new therapeutic opportunities. Cell Death and Differentiation vol. 28 423–426 Preprint at 10.1038/s41418-020-00703-w (2021).

19. Frescas, D. & Pagano, M. Deregulated proteolysis by the F-box proteins SKP2 and β-TrCP: Tipping the scales of cancer. Nature Reviews Cancer vol. 8 438–449 Preprint at 10.1038/nrc2396 (2008).

20. Lau, A. W., Fukushima, H. & Wei, W. The Fbw7 and Beta-TRCP E3 ubiquitin ligases and their roles in tumorigenesis. Frontiers in Bioscience vol. 17 2197–2212 Preprint at 10.2741/4045 (2012).

21. Bi, Y., Cui, D., Xiong, X. & Zhao, Y. The characteristics and roles of β-TrCP1/2 in carcinogenesis. FEBS Journal vol. 288 3351–3374 Preprint at 10.1111/febs.15585 (2021).

22. Herrero-Ruiz, J. et al. βTrCP controls the lysosome-mediated degradation of CDK1, whose accumulation correlates with tumor malignancy. Oncotarget 5, 7563–7574 (2014).

23. Hart, M. et al. The F-box protein β-TrCP associates with phosphorylated β-catenin and regulates its activity in the cell. Current Biology 9, 207–211 (1999).

24. Jaffry, U. & Wells, G. Small molecule and peptide inhibitors of βTrCP and the βTrCP–NRF2 protein–protein interaction. Biochemical Society Transactions vol. 51 925–936 Preprint at 10.1042/BST20220352 (2023).

25. Limón-Mortés, M. C. et al. UV-induced degradation of securin is mediated by SKP1-CUL1-βTrCP E3 ubiquitin ligase. J Cell Sci 121, 1825–1831 (2008).

26. Giráldez, S. et al. G1/S phase progression is regulated by PLK1 degradation through the CDK1/βTrCP axis. FASEB Journal 31, 2925–2936 (2017).

27. Piscitello, D. et al. AKT overactivation can suppress DNA repair via p70S6 kinase-dependent downregulation of MRE11. Oncogene 37, 427–438 (2018).

28. Robertson, H., Hayes, J. D. & Sutherland, C. A partnership with the proteasome; the destructive nature of GSK3. Biochemical Pharmacology vol. 147 77–92 Preprint at 10.1016/j.bcp.2017.10.016 (2018).

29. Pearce, L. R. et al. Characterization of PF-4708671, a novel and highly specific inhibitor of p70 ribosomal S6 kinase (S6K1). Biochemical Journal 431, 245–255 (2010).

30. Stambolic, V., Ruel, L. & Woodgett, J. R. Lithium inhibits glycogen synthase kinase-3 activity and mimics wingless signalling in intact cells. Current Biology 6, 1664–1669 (1996).

31. Bennett, C. N. et al. Regulation of Wnt signaling during adipogenesis. Journal of Biological Chemistry 277, 30998–31004 (2002).

32. Watanabe, N. et al. Cyclin-dependent kinase (CDK) phosphorylation destabilizes somatic Wee1 via multiple pathways. Proc Natl Acad Sci U S A 102, 11663–11668 (2005).

33. Belmonte-Fernández, A. et al. Cisplatin-induced cell death increases the degradation of the MRE11-RAD50-NBS1 complex through the autophagy/lysosomal pathway. Cell Death Differ 30, 488–499 (2023).

34. Stracker, T. H., Couto, S. S., Cordon-Cardo, C., Matos, T. & Petrini, J. H. J. Chk2 Suppresses the Oncogenic Potential of DNA Replication-Associated DNA Damage. Mol Cell 31, 21–32 (2008).

35. Barbour, H. et al. An inventory of crosstalk between ubiquitination and other post-translational modifications in orchestrating cellular processes. iScience vol. 26 106276 Preprint at 10.1016/j.isci.2023.106276 (2023).

36. Chapman, J. R. & Jackson, S. P. Phospho-dependent interactions between NBS1 and MDC1 mediate chromatin retention of the MRN complex at sites of DNA damage. EMBO Rep 9, 795–801 (2008).

37. Wang, Q. et al. Rad17 recruits the MRE11-RAD50-NBS1 complex to regulate the cellular response to DNA double-strand breaks. EMBO J 33, 862–877 (2014).

38. Wu, J. et al. Skp2 E3 Ligase Integrates ATM Activation and Homologous Recombination Repair by Ubiquitinating NBS1. Mol Cell 46, 351–361 (2012).

39. Kim, H., Kim, D., Choi, H., Shin, G. & Lee, J. K. Deubiquitinase USP2 stabilizes the MRE11–RAD50–NBS1 complex at DNA double-strand break sites by counteracting the ubiquitination of NBS1. Journal of Biological Chemistry 299, (2023).

40. Ha, G. H. et al. Pellino1 regulates reversible ATM activation via NBS1 ubiquitination at DNA double-strand breaks. Nat Commun 10, (2019).

41. Hodge, C. D. et al. RNF8 E3 ubiquitin ligase stimulates Ubc13 E2 conjugating activity that is essential for DNA double strand break signaling and BRCA1 tumor suppressor recruitment. Journal of Biological Chemistry 291, 9396–9410 (2016).

42. Nowsheen, S. et al. L3MBTL2 orchestrates ubiquitin signalling by dictating the sequential recruitment of RNF8 and RNF168 after DNA damage. Nat Cell Biol 20, 455–464 (2018).

43. Escribano-Díaz, C. et al. A Cell Cycle-Dependent Regulatory Circuit Composed of 53BP1-RIF1 and BRCA1-CtIP Controls DNA Repair Pathway Choice. Mol Cell 49, 872–883 (2013).

44. Jin, J. et al. SCFβ-TRCP links Chk1 signaling to degradation of the Cdc25A protein phosphatase. Genes Dev 17, 3062–3074 (2003).

45. Busino, L. et al. Degradation of Cdc25A by β-TrCP during S phase and in response to DNA damage. Nature 426, 87–91 (2003).

46. Inuzuka, H. et al. Phosphorylation by Casein Kinase I promotes the turnover of the Mdm2 oncoprotein via the SCFβ-TRCP ubiquitin ligase. Cancer Cell 18, 147–159 (2010).

47. Watanabe, N. et al. M-phase kinases induce phospho-dependent ubiquitination of somatic Wee1 by SCFβ-TrCP. Proc Natl Acad Sci U S A 101, 4419–4424 (2004).

48. Mailand, N., Bekker-Jensen, S., Bartek, J. & Lukas, J. Destruction of Claspin by SCFβTrCP Restrains Chk1 Activation and Facilitates Recovery from Genotoxic Stress. Mol Cell 23, 307–318 (2006).

49. Saitoh, T. & Katoh, M. Expression profiles of βTRCP1 and βTRCP2, and mutation analysis of βTRCP2 in gastric cancer. Int J Oncol 18, 959–964 (2001).

50. KIM, C. J. et al. Somatic mutations of the β-TrCP gene in gastric cancer. APMIS 115, 127–133 (2007).

51. Gerstein, A. V. et al. APC/CTNNB1 (β-catenin) pathway alterations in human prostate cancers. Genes Chromosomes Cancer 34, 9–16 (2002).

52. Wood, L. D. et al. The Genomic Landscapes of Human Breast and Colorectal Cancers. Science (1979) 318, 1108–1113 (2007).

53. Turgeon, M. O., Perry, N. J. S. & Poulogiannis, G. DNA damage, repair, and cancer metabolism. Frontiers in Oncology vol. 8 342110 Preprint at 10.3389/fonc.2018.00015 (2018).

54. Sobanski, T. et al. Cell Metabolism and DNA Repair Pathways: Implications for Cancer Therapy. Frontiers in Cell and Developmental Biology vol. 9 633305 Preprint at 10.3389/fcell.2021.633305 (2021).

55. Margottin, F. et al. A novel human WD protein, h-βTrCP, that interacts with HIV-1 Vpu connects CD4 to the ER degradation pathway through an F-box motif. Mol Cell 1, 565–574 (1998).

56. Galindo-Moreno, M. et al. SCF(FBXW7)-mediated degradation of p53 promotes cell recovery after UV-induced DNA damage. FASEB Journal 33, (2019).

57. Giráldez, S. et al. SCFFBXW7α modulates the intra-S-phase DNA-damage checkpoint by regulating Polo like kinase-1 stability. Oncotarget 5, 4370–4383 (2014).

58. Mancinelli, L. et al. A class of DNA-binding peptides from wheat bud causes growth inhibition, G2 cell cycle arrest and apoptosis induction in HeLa cells. Mol Cancer 8, 1–11 (2009).

